# Functional Antagonism Between CELF and Mbnl Proteins in the Cytoplasm

**DOI:** 10.1101/009183

**Authors:** Eric T. Wang, Amanda J. Ward, Jennifer Cherone, Thomas T. Wang, Jimena Giudice, Thomas A. Cooper, Christopher B. Burge

## Abstract

The conserved CUGBP1, Elav-like (CELF) family of RNA binding proteins contribute to heart and skeletal muscle development and are implicated in myotonic dystrophy (DM). To understand genome-wide functions of CELF proteins, we analyzed transcriptome dynamics following induction of *CELF1* or *CELF2* in adult mouse heart or *CELF1* in muscle by RNA-seq, complemented by crosslinking/immunoprecipitation-sequencing (CLIP-seq) analysis of mouse cells and tissues to distinguish direct from indirect regulatory targets. Analysis of expression and mRNA binding data revealed hundreds of mRNAs bound in their 3' UTRs by both CELF1 and and the developmentally induced Mbnl1 protein, 3-fold more than expected. The relative extent of CELF1 and Mbnl1 binding in 3' UTRs predicted the extent of repression or stabilization, respectively, following CELF induction. These findings support a “Cytoplasmic Competition” model in which CELF and Mbnl proteins compete to specify degradation or membrane localization/stabilization, respectively, of an overlapping set of targets. Several hundred messages contained proximal CELF1 and Mbnl1 binding sites (within 50 bases), and were more strongly repressed by CELF1 than messages with distal sites. Messages with different spacing of CELF and Mbnl sites in their 3' UTRs exhibited different developmental dynamics, suggesting that spacing is used to tune cytoplasmic competition between these factors to specify the timing of developmental induction. CELF1 also shared dozens of splicing targets with Mbnl1, most regulated oppositely, confirming a phenomenon observed in smaller scale studies but not previously supported by genome-wide methods, which also appears to enhance developmental transitions.

## Introduction

The CUGBP1, Elav-like (CELF) family of conserved and developmentally regulated RNA binding proteins play roles in early embryonic development, heart and skeletal muscle function, and have been implicated in myotonic dystrophy (DM) (Timchenko et al., 1996), and suggested to contribute to other diseases (Ladd, 2013). The six family members expressed in mammals can be divided into two subfamilies: CELF1/CELF2, which are expressed most highly in heart, skeletal muscle, and brain, and CELF 3-6 which exhibit more restricted expression, with *CELF3* and *CELF5* in the brain, and *CELF6* in the brain, kidney and testis. The CELF proteins contain two N-terminal RNA recognition motifs (RRM) and one C-terminal RRM, with which they bind their targets, and a linker region termed the ‘divergent domain’ separating RRM2 and RRM3 that is involved in regulation of alternative splicing and mRNA decay (Han and Cooper, 2005; Vlasova et al., 2008).

During normal development, CELF 1 and CELF2 proteins are highly expressed in early embryonic stages, and are then downregulated more than 10-fold in skeletal muscle (Ladd et al., 2005) and heart (Kalsotra et al., 2008) during postnatal development, remaining at low levels in adult tissues. This developmental downregulation occurs through multiple mechanisms, including repression by microRNAs (miRNAs) and destabilizing reductions in protein phosphorylation (Kalsotra et al., 2014; Kalsotra et al., 2010). However, in DM type 1 (DM1), CELF1 protein levels increase in heart and skeletal muscle (Timchenko et al., 2004) as the protein is stabilized by PKC-mediated phosphorylation (Kuyumcu-Martinez et al., 2007).

The combination of increased CELF levels and MBNL sequestration by CUG repeat RNA is thought to be responsible for much of DM pathology by reversing the developmental changes in both proteins toward embryonic levels, shifting splicing of regulatory targets toward fetal isoforms (Ho et al., 2004; Kuyumcu-Martinez et al., 2007; Lin et al., 2006; Philips et al., 1998). CELF1 re-expression in adults recapitulates a subset of the mis-regulated splicing events observed in DM1 skeletal muscle and heart (Kalsotra et al., 2010; Kalsotra et al., 2008; Ward et al., 2010). Out of 44 developmentally regulated alternative splicing events in heart development that were investigated, 24 were found to revert toward embryonic splicing levels in response to inducible expression of *CELF1* in the adult heart or in mice lacking *Mbnl1* (Kalsotra et al., 2008). A long-standing question has been whether these splicing factors regulate a shared set of exons, and whether they do so in an antagonistic fashion. The half dozen events known to be regulated by both CELF and Mbnl proteins, including *H2afy* exon 6 and *Mbnl2* exon 8, are regulated antagonistically (Dhaenens et al., 2008; Ho et al., 2005; Kalsotra et al., 2008). However, in a study analyzing CELF motifs present near Mbnl-responsive alternative exons, evidence for widespread antagonistic regulation was not observed (Du et al., 2010).

CELF proteins are present in both cytoplasm and nucleus and have been shown to play roles in deadenylation, RNA stability and translation as well as splicing (Paillard et al., 2003; Timchenko et al., 2004; Vlasova et al., 2008). Human CELF1 is 88% identical to the *Xenopus* homolog Embryo Deadenylation Element binding protein (EDEN-BP) and can functionally replace it, binding to the EDEN to control polyA tail length and aiding in rapid deadenylation of maternal mRNAs after fertilization (Paillard et al., 2003). CELF1 interacts with polyA-specific ribonuclease (PARN) in HeLa cell extracts to promote deadenylation of *C-FOS* and *TNFalpha* transcripts (Moraes et al., 2006). Furthermore, siRNA-mediated knockdown of *CELF1* in HeLa cells and myoblasts led to the stabilization of a set of normally rapidly degraded transcripts bound by CELF at GU-rich elements (GRE) and GU-repeats (Lee et al., 2010; Rattenbacher et al., 2010; Vlasova et al., 2008). It has recently been established that many genes shift toward greater expression of longer 3' UTR isoforms during differentiation of muscle cells and likely other cell types (Ji et al., 2009). It is possible that upregulation of CELF proteins in adult tissues could affect the stability of many transcripts and contribute to DM pathology.

To investigate the functional significance of the postnatal downregulation of CELF in heart and skeletal muscle tissues, we inducibly expressed *CELF1* in adult mouse heart or muscle or *CELF2* in adult heart and performed a time series RNA-seq analysis following induction. To distinguish direct from indirect targets and determine rules relating binding to regulation, we also identified transcriptome-wide binding locations by crosslinking/immunoprecipitation-sequencing (CLIP-seq) analysis of mouse cells and tissues. Our analysis of splicing identified hundreds of direct CELF targets, showing a strong inverse relationship with heart developmental changes and a tendency to counteract the effects of Mbnl on splicing, confirming a phenomenon observed in smaller scale studies but not previously supported by genome-wide methods.

Our analysis of 3' UTR binding revealed a number of new phenomena, including 3-fold more mRNAs bound by both CELF and Mbnl proteins than expected, nearby spacing of CELF and Mbnl sites in mRNAs, and that relative abundance of CELF and Mbnl sites in 3' UTRs predicts the extent of CELF-directed downregulation or stabilization, respectively, in a manner that depends on site spacing. Together, these observations lead to a Cytoplasmic Competition model in which cytoplasmic CELF and Mbnl proteins compete to specify different mRNA fates to tune the expression and localization of mRNAs during development. Our approach, based on transcriptome-wide analysis of binding and isoform-level expression, complements previous molecular genetic and biochemical analyses of these factors, by emphasizing general trends that impact dozens or hundreds of messages.

## Results

### Identification of Hundreds of CELF-Responsive Exons

We performed strand-specific paired-end RNA sequencing (RNA-seq) of poly(A)+ RNA from skeletal muscle or heart of mice in which *CELF1* was induced, or heart of mice in which *CELF2* was induced at several time points post-induction, and from control mice, in biological triplicate. The *CELF1* mice have been described previously (Koshelev et al., 2010; Ward et al., 2010), the *CELF2* mouse model is described in Methods, and all mice used are summarized in Supplemental Table S1. Human *CELF* transgenes were induced in mice by administration of doxycycline, in separate mouse lines for each tissue/protein pair, and reached levels between 5-and 10-fold above endogenous levels (Fig. S1a,b). The mRNA level of CELF1 declined at the end of the time course, likely as a secondary effect of heart pathology on the alpha MHC promoter that drives rtTA (Lowes et al., 1997). However, CELF1 protein levels consistently increase in these mice at least 8-fold in skeletal muscle (Ward et al., 2010) and ∼4-fold in heart and remain high through at least day 7 (Koshelev et al., 2010). Heart pathology was not observed in the *CELF2* mouse, and CELF2 RNA and protein levels increased throughout the time course (Fig. S1c,d). RNA was isolated from three mice each at 12 hours (h), 24 h, 72 h, and 7 days (d) following induction, and three control mice lacking tet-inducible *CELF1* or *CELF2* transgenes treated with doxycycline for 72 h. MISO was used to estimate percent spliced in (PSI or Ψ) values for cassette (skipped) exons, alternative 5' and 3' splice sites, and retained introns. For each comparison of Ψ between samples, we calculated a Bayes Factor (BF), representing the ratio of the likelihood of the hypothesis that PSI values differ relative to the null hypothesis of no change in PSI, and used BFs to identify differentially regulated exons.

We identified thousands of exons whose Ψ values changed between time points at a BF cutoff of 5. For example, exon 3 in Tmed2 had a Ψ of 79% in control heart, but decreased upon *CELF1* induction to 13% at seven days (Fig. 1a). To identify splicing events that change monotonically over time, we developed a permutation-based method that could be applied to time course data. We ordered samples chronologically, and for each event compared all pairs of samples from different time points, tallying the number of comparisons representing significant increase or significant decrease in Ψ (at BF > 5). We calculated a quantity called δ, the number of significant positive ΔΨ values (increases over time) minus the number of significant negative ΔΨ values (Fig. S2a). To assess statistical significance, we recalculated δ after randomly permuting the sample labels. Repeating this process 100 times, we generated a null distribution, and derived a “monotonicity Z-score” (MZ = (δ − µ) / σ), where µ and σ are the mean and standard deviation of the null distribution, respectively. Splicing events with large MZ values change consistently over the time course (MZ-scores and Ψ values for all experiments are listed in Table S2). We observed 627 and 825 exons responsive to *CELF1* induction at a MZ score of 1.8 in heart and skeletal muscle, respectively.

**Figure 1.**
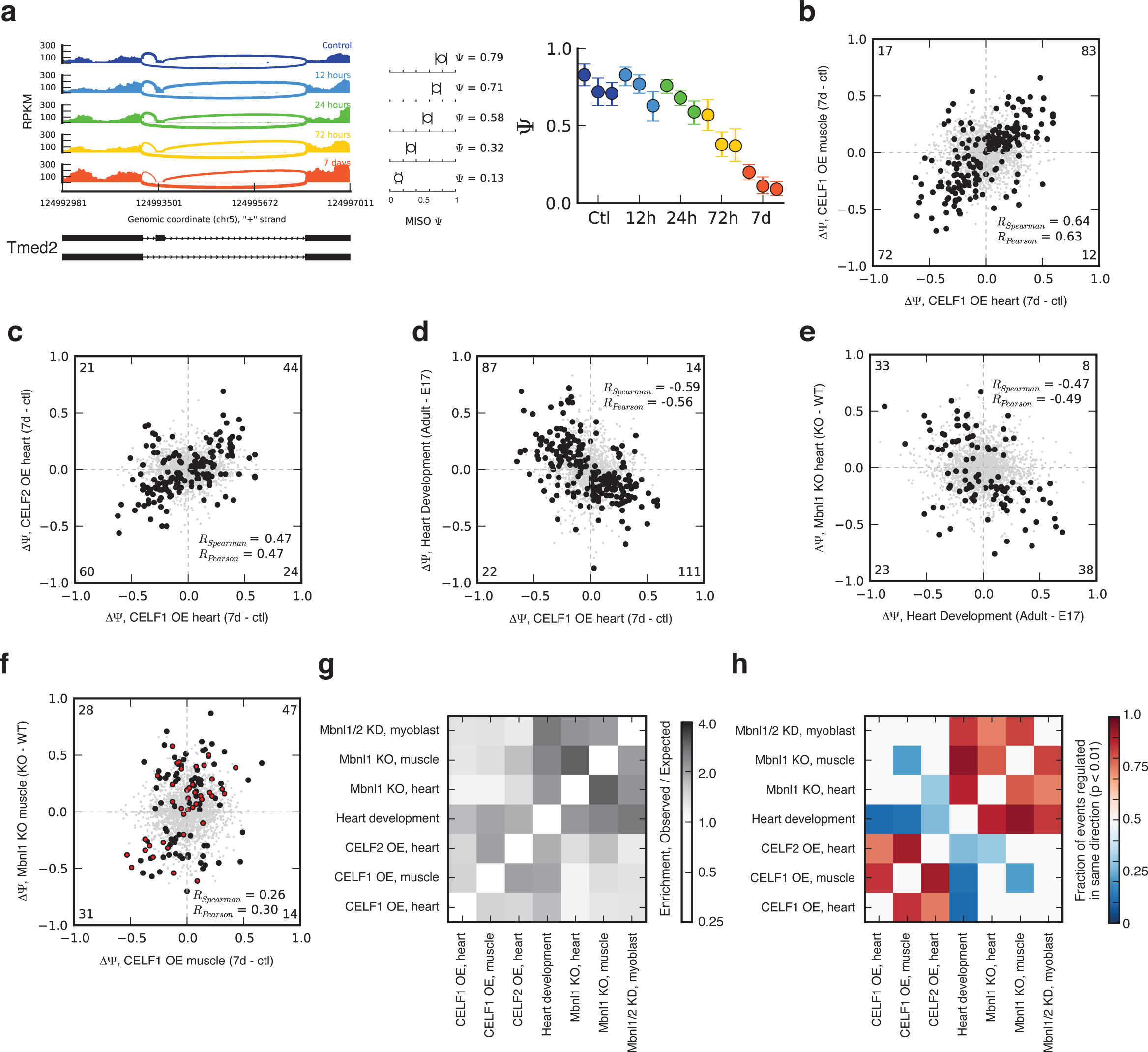
CELF1 and CELF2 regulate hundreds of splicing events, reversing many changes during heart development, and antagonizing a subset of Mbnl-regulated exons. a) RNA-seq read coverage across Tmed2 exon 3 from mouse heart at several time points following CELF1 induction. MISO Ψ values and 95% confidence intervals shown at right. b) Splicing changes that occur following CELF1 over-expression in heart correlate with splicing changes that occur following CELF1 over-expression in muscle. *n* = 2496 alternative splicing events (skipped exons, alternative 3' splice sites, alternative 5' splice sites, retained introns, and mutually exclusive exons) shown: monotonically changing exons shown as black circles, others as gray dots. Correlation values of monotonic events (shown) are higher than for all events: the numbers of monotonic events in each quadrant are shown in corners. c) Splicing changes that occur in response to CELF1 over-expression in heart correlate with splicing changes that occur in response to CELF2 over-expression in heart. As in c) with *n* = 2129 skipped exons shown. d) Splicing changes that occur in response CELF1 over-expression in heart inversely correlate with splicing changes that occur during mouse heart development. As in c) with *n* = 1952 skipped exons shown. e) Splicing changes that occur in Mbnl1 KO heart inversely correlate with splicing changes that occur during mouse heart development. *n* = 3190 skipped exons shown, as in Fig. 1c. f) Splicing changes that occur in Mbnl1 KO muscle correlate with splicing changes that occur following CELF1 over-expression in muscle. *n* = 1501 skipped exons shown. Events that changed monotonically, or with BF > 5, following CELF1 over-expression in muscle or Mbnl1 KO in muscle, respectively, are shown in black, and correlations are listed for these events. Events that also changed monotonically during heart development are shown in red. g) Exons regulated during heart development also tend to change in Mbnl1 and Mbnl2 knockdown myoblasts, and in Mbnl1 and Mbnl2 knockout mice, and CELF1 or CELF2 over-expressing mice. Enrichment (observed / expected number of regulated exons) is shown in the heatmap. h) Splicing of exons regulated in response to CELF over-expression or Mbnl depletion tends to change in a direction opposite to changes that occur during heart development. The fraction of events changing in the same direction for each pair of comparisons is shown in the heat map (only biases significant at P < 0.01 by binomial test are colored). Also see Figures S1-2 and Tables S1-3.

### About 30% of splicing changes in heart development respond to *CELF1* induction

We next sought to understand the functions of CELF proteins in different tissues and developmental stages. Comparing ΔΨ values of monotonically changing skipped exons between *CELF1*-induced heart and *CELF1*-induced skeletal muscle, we observed a fairly high correlation (R_Spearman_ = 0.64), suggesting that the functions and regulatory targets of CELF1 in these tissues are quite similar (Fig. 1b). To ask about functions of different CELF family members, we compared splicing changes in heart following induction of *CELF1* or *CELF2*. This comparison yielded a moderate positive correlation (R_Sp_ = 0.47), suggesting similar but not identical splicing functions (Fig. 1c).

To explore connections to developmental roles of CELF proteins, we compared splicing in *CELF1* heart to changes that occur during normal heart development. Developmental changes were analyzed by use of an RNA-seq time series including embryonic day 17 (E17), postnatal day (PN) 1, PN 10, PN 28, and adult, during which CELF1 protein levels fall by > 10-fold (Kalsotra et al., 2008). In all, 234 skipped exons changed monotonically in heart development and/or *CELF1* induction. Notably, splicing of 198 (85%) of these exons changed in the opposite direction following *CELF1* induction as they did during heart development (R_Sp_ = −0.59, Fig. 1d). This widespread reversion of developmental changes suggests that normal reductions in CELF activity during heart development may contribute to a substantial portion – perhaps 30% (Fig. S2b) – of the splicing changes that occur during normal heart development (Giudice et al., 2014; Kalsotra et al., 2008).

### A large cohort of exons is regulated antagonistically by CELF and Mbnl proteins in the nucleus

Six mouse exons (Kalsotra et al., 2008) and at least one human exon (Ho et al., 2004) are known to be regulated antagonistically by CELF1 and Mbnl1. Confirming and extending these observations, we identified dozens of additional exons regulated by both factors. Mbnl1 depletion (Wang et al., 2012) mimicked a reversal of many splicing changes that occur during heart development (R_Sp_ = –0.47, Fig. 1e). In skeletal muscle, we detected 120 exons whose splicing was responsive to both proteins, of which 78 (65%) were regulated in an antagonistic fashion (Fig. 1f; exons responsive to both factors are listed in Table S3). Extending this analysis to additional tissues and cell lines, and comparing to heart development, we found that exons responsive to induction of CELF1 or CELF2 showed a strong and quite general tendency to also respond to Mbnl depletion and heart development (Fig. 1g). Furthermore, the direction of splicing regulation tended to reverse developmental patterns, with both Mbnl depletion and CELF induction often mimicking reversal of developmental changes (Fig. 1h, Table S3). The 206 total splicing events responsive to both Mbnl1 depletion and CELF1 re-expression in muscle and/or heart were enriched for particular Gene Ontology categories, including “cell differentiation”, “multicellular organismal development”, “microtubule cytoskeleton”, and “cell junction”, suggesting roles in developmental remodeling of the heart. Furthermore, 124 events (60%) were reading frame-preserving (compared to background of 44%, p < 0.001, binomial test), suggesting that CELF and Mbnl proteins play a substantial role in shaping the heart proteome. These observations provide genome-wide evidence for the principle that CELF and Mbnl proteins exert opposing effects on the splicing of a large cohort of exons. They also expand the number of developmentally regulated CELF-and Mbnl-responsive exons several-fold.

### Transcriptome-wide binding locations of CELF1 in heart, muscle, and myoblasts

To identify transcript sites bound by CELF1, we performed CLIP-seq analysis using the 3B1 antibody against the endogenous protein, yielding 1.6 million (M), 1.0 M, and 1.6 M reads uniquely mapping to the genome and splice junctions in 16 week-old C57BL/6 heart, 16 week-old C57/BL6 muscle, and C2C12 myoblast samples, respectively, after collapsing identical reads. Mapping predominantly occurred to transcribed regions, with enrichment for introns and/or 3' UTRs (Fig. S3a).

CELF1 binding locations were consistently observed across different tissues, for example in the 3' UTR of the myeloid-associated differentiation marker gene, *Myadm* (Fig. 2a). CELF1 binding density along mRNAs, assessed “locally” using 5 nucleotide (nt) windows, was highly correlated across tissues and cell lines, and distinct from that observed for Mbnl1 (Fig. 2b). Controlling for pre-mRNA length and gene expression, CELF1 CLIP clusters were enriched for UGU-containing pentanucleotides (5mers) in heart and other tissues (Fig. 2c, Fig. S3b). Introns flanking alternative exons with CLIP clusters within 1 kb from either splice site were more highly conserved across species, in particular in the downstream intron, supporting their in vivo function (Fig. 2d).

**Figure 2.**
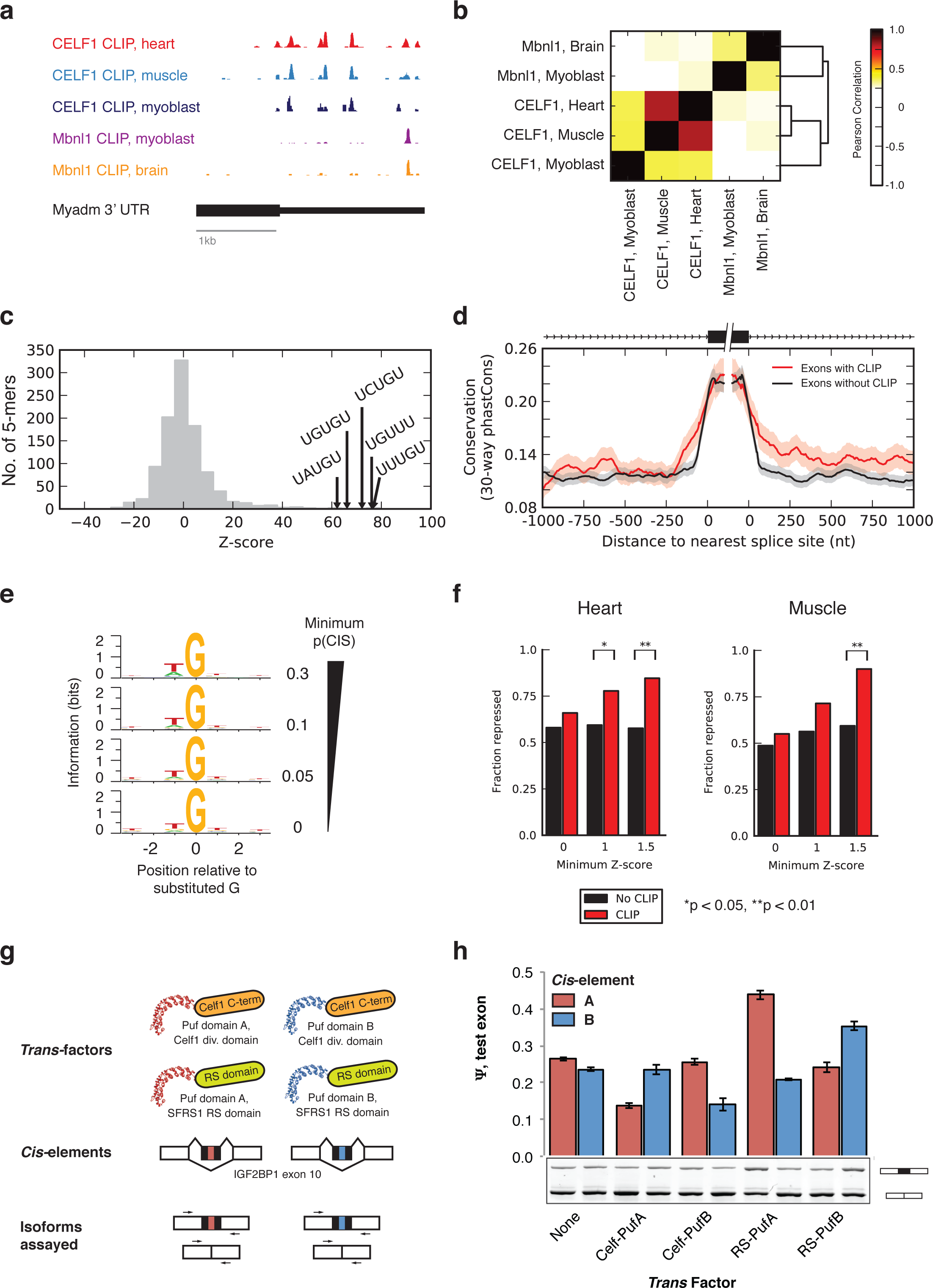
CELF1 binds to consistent locations across cell and tissues, and represses splicing of bound exons. a) CELF1 CLIP-seq read coverage across the 3' UTR of mouse *Myadm* gene. Binding locations are highly correlated between muscle and heart. b) Correlation of CLIP tag densities in 5 nt windows across all 3' UTRs expressed in mouse heart, muscle, and myoblasts. c) Histogram of enrichment Z-scores of 5mers based on frequency of occurrence in CELF1 heart CLIP clusters relative to control regions from the 3' UTRs. d) Meta-exon analysis of conservation (mean + 95% confidence interval of phastCons score in 5 nt windows shown) at a range of distances from 3' and 5' splice sites of exons with (red) and without (black) overlapping CELF1 CLIP clusters in heart. n = 432 skipped exons shown. e) Information content (relative entropy compared to uniform) of genomic positions in regions where CELF1 CLIP-seq reads map, grouped by the frequency of substitution in CLIP reads relative to genome of the central G position. f) Fraction of significantly repressed exons (Ψ at 7 days < Ψ in control animals) with or without CLIP clusters, at three monotonicity Z-score thresholds from CELF1 over-expression time courses in heart and muscle. g) Design of experiment involving splicing reporters and Pumilio-based synthetic splicing factors, and assessment of splicing by qRT-PCR. h) Tethering the non-RRM regions of CELF1 to a cassette exon in a splicing reporter by Pumilio fusion leads to exon skipping. In a positive control, tethering of RS domains to the cassette exon enhances exon inclusion. Enhancement by RS and repression by CELF1 occur only when the pumilio domain has affinity for the inserted cis-element. Also see Figures S3-4.

To more precisely map sites of CELF1 binding, we measured the frequency of crosslink-induced substitutions (CIS) – positions where CLIP-seq reads differ from the genome (Kishore et al., 2011; Wang et al., 2012) – at each position within CLIP clusters. We noted that guanosines with high CIS showed biases in flanking bases, with the –1 base preceding the substituted guanosine increasingly biased towards uracil as CIS frequency increased (reaching ∼61% in heart), and the base at the +1 position was also biased toward uracil (Fig. 2e and Fig. S3c). The implied CIS-enriched motif, UGU, resembles the motifs observed in Figure 2c and in previous studies of CELF1 binding affinity (Lambert et al., 2014; Lambert, 2014; Marquis et al., 2006), suggesting that frequently substituted guanosines are highly enriched for sites of direct crosslinking to CELF1 protein.

### Context-dependent regulation of splicing by CELF1

Exons whose splicing changed after *CELF1* induction were enriched for CELF1 CLIP clusters (Fig. S4). To identify an RNA map describing the splicing regulatory activity of CELF1, we analyzed the location of CELF1 clusters relative to exons responsive to *CELF1* over-expression in heart and/or muscle. We found that cassette exons bound by CELF1 were more likely to be repressed following CELF1 induction in heart and muscle (Fig. 2f), with 80-85% of exons with MZ score > 1.5 being repressed. Though we had reasonable statistical power, we did not observe consistent repression or activation of exons bound by CELF1 in the upstream or downstream introns. These observations suggest that exonic binding by CELF family members may directly repress splicing, while the direction of splicing regulation resulting from intronic binding may depend on other variables such as RNA structure or cooperation or antagonism with other RNA binding proteins (Dembowski and Grabowski, 2009; Goo and Cooper, 2009).

To directly test the ability of CELF1 to repress splicing when bound to a cassette exon, we fused the C-terminus of CELF1 (omitting the RRM domains) to two different Pumilio (Puf) domains, each recognizing a distinct 8-base Puf motif (Wang et al., 2009a). We expressed these constructs in Hea cells, along with a splicing reporter (Xiao et al., 2009) containing either cognate or non-cognate Puf motifs within the cassette exon, and used quantitative RT-PCR to assess splicing changes (Fig. 2g). We found that expression of CELF1-Puf fusion decreased Ψ from ∼25% to ∼12% when paired with the cognate Puf motif, but had negligible effect when paired with the non-cognate motif. Thus, recruitment of a single CELF1 protein to an exon appears sufficient to reduce splicing by double digit percent values in this system. Natural targets often have multiple CELF1 binding sites, potentially magnifying the impact on splicing, as is commonly observed for splicing regulatory factors (Matlin et al. 2005). Control RS domain-Puf fusions activated splicing, as expected (Fig. 2h) (Wang et al., 2009a). These observations indicate that tethering the CELF1 C-terminus to an exon is sufficient to repress its splicing.

### Dose-dependent mRNA Decay Associated with CELF1 binding to 3' UTRs

We observed extensive CELF1 binding to 3' UTRs, with increased density upstream of the cleavage and polyadenylation site (PAS) (Fig. 3a). Consistent with the ability of CELF1 to recruit cytoplasmic deadenylases (Vlasova-St Louis et al., 2013), binding density increased close to the PAS on the upstream side and fell to background levels just downstream of the PAS site in heart, muscle, and myoblast. Increased phylogenetic conservation was observed for 3' UTRs containing CELF1 CLIP clusters, relative to 3' UTRs with similar length and expression level, suggesting that these mRNAs are enriched for conserved functional motifs or structures (Fig. 3b). Gene ontology analysis of 3' UTR targets of CELF1 revealed enrichment of proteins related to muscle structures such as M band and I band, and factors involved in vesicle and protein transport, in both heart and muscle (data not shown).

**Figure 3.**
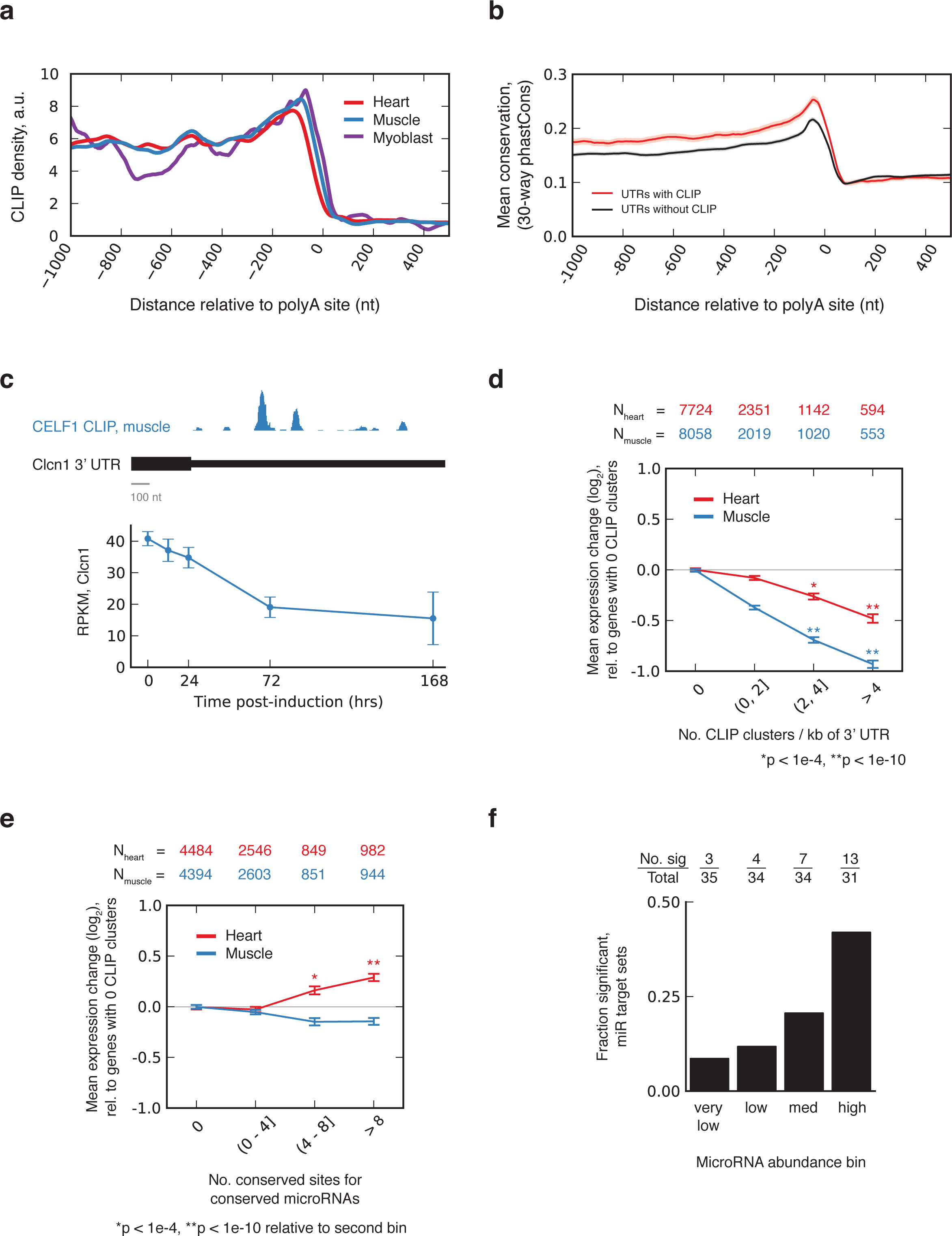
CELF1 binds to 3' UTRs and regulates message stability in a dose-dependent fashion. a) Mean CELF1 CLIP density at positions along 3' UTRs in heart, muscle, and myoblasts. b) Mean conservation in sets of 3' UTRs with and without CELF1 CLIP clusters with similar expression levels and UTR lengths (shading represents SEM). c) Expression of Clcn1 in muscle based on RNA-seq (mean ± standard deviation), at various times following CELF1 overexpression (below); CLIP density in *Clcn1* 3' UTR (above). d) Mean log expression change following CELF1 induction (7 d over control) for transcripts grouped by number of CELF1 CLIP clusters in their 3' UTRs. Transcripts with greater CLIP cluster density are down-regulated more strongly in heart and muscle (number of genes in each category listed above). e) As in d), but for genes grouped by number of conserved target sites to conserved miRNAs in their 3' UTRs. Transcripts with greater numbers of conserved miRNA seed matches are de-repressed in heart but not in muscle. f) The expression changes of miRNA target gene sets was analyzed, and the number of sets with significant derepression (P < 1e-4 by modified KS test) is listed above for each of four cohorts of miRNAs, grouped by miRNA abundance in heart, from lowest to highest. Target sets corresponding to highly expressed miRNAs are more often derepressed. Also see Table S4.

To explore potential effects of CELF1 on mRNA stability, we studied changes in expression of CELF1-bound mRNAs following CELF1 induction. For example, the *Clcn1* mRNA, which had large CELF1 binding clusters in its 3' UTR, decreased in expression 50-70% from its initial level 7 days after *CELF1* induction, in both heart and muscle (Fig. 3c). Analyzing gene expression globally, we found that mean expression of CELF1-bound genes decreased substantially at 7 days post-induction relative to genes not bound by CELF1 (Table S4). Furthermore, the mean extent of target down-regulation increased monotonically with the density of CLIP clusters in the 3' UTR to 1.4-fold and 2-fold in heart and muscle, respectively, for messages with 4 or more CLIP clusters per kb of 3' UTR (Fig. 3d).

We expected to see similar levels of CELF1-mediated target down-regulation in heart and muscle, given that the level of *CELF1* induction was similar in both tissues (Fig. S1) (Koshelev et al., 2010; Ward et al., 2010). However, the magnitude of target downregulation in heart was consistently only about half of that observed in muscle for messages with similar CLIP density. Considering additional variables that might impact the magnitude of regulation, we noticed that genes with numerous seed matches to conserved miRNAs were de-repressed relative to genes without miRNA sites in the heart; however, this pattern was not observed in muscle (Fig. 3e). Furthermore, when we grouped miRNAs by their expression level within heart tissue, we found that the most strongly de-repressed mRNAs tended to have seed matches to the highest expressed heart miRNAs (Fig. 3f). Though further investigation is required, these observations hint that, in heart, CELF1 induction may somehow lead to decreased miRNA activity, partially mitigating the impact of CELF1-directed mRNA decay.

### Antagonistic mRNA regulation by CELF and Mbnl Proteins in the cytoplasm

In addition to their nuclear functions in RNA splicing, Mbnl proteins also exert cytoplasmic functions, targeting 3' UTR-bound mRNAs for localization to membrane destinations and promoting translation of targeted messages (Adereth et al., 2005; Wang et al., 2012). To ask whether this function might be related to cytoplasmic functions of CELF, we compared CELF1 and Mbnl1 3' UTR binding targets. Strikingly, we observed over a thousand 3' UTRs bound by both CELF1 and Mbnl1, three times as many as expected by chance, controlling for expression (Fig. 4a; CLIP cluster coordinates are listed in Table S5). This observation led us to explore whether CELF and Mbnl might exert antagonistic, or synergistic, effects on mRNAs in the cytoplasm.

**Figure 4.**
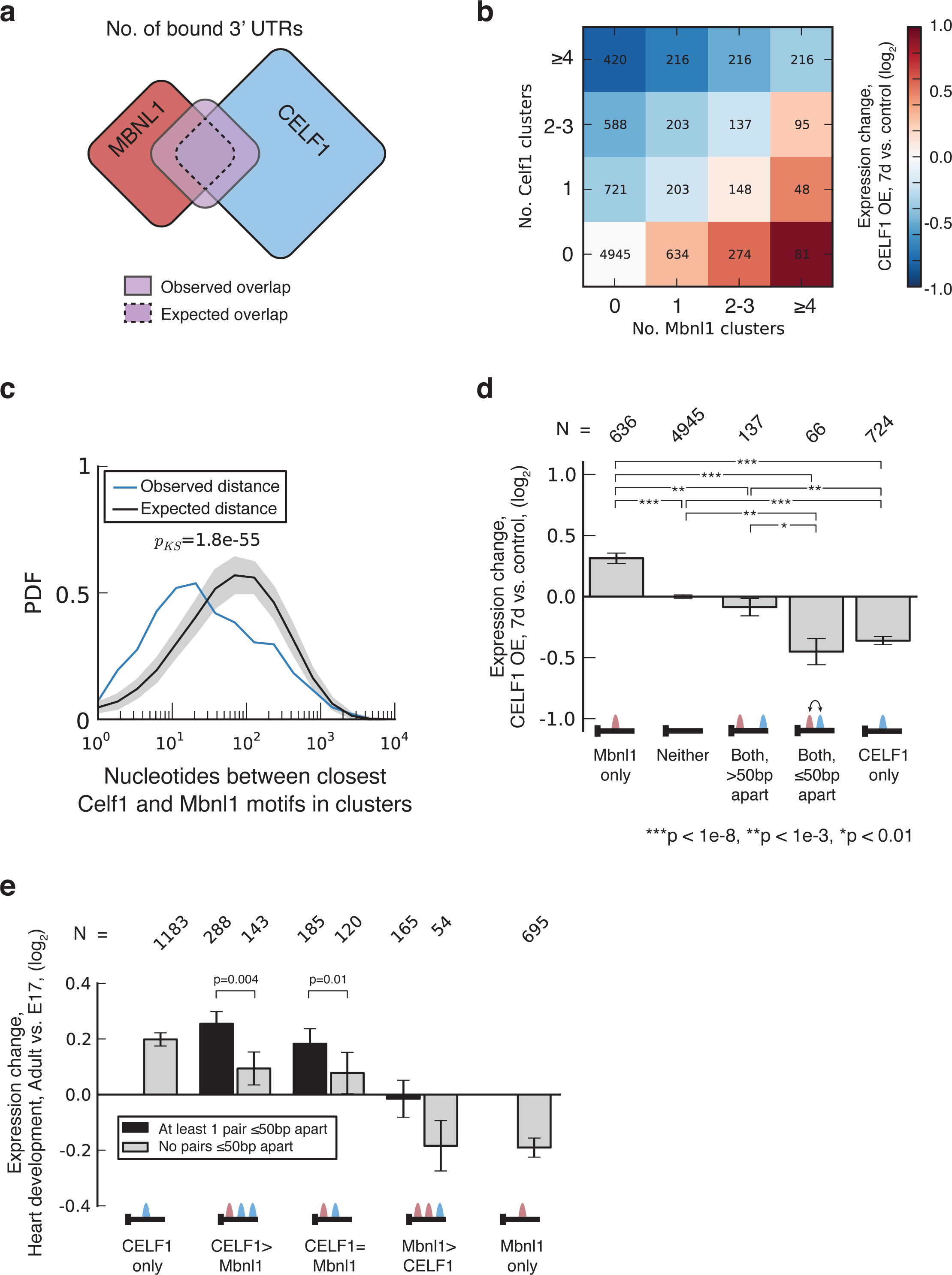
CELF1 and Mbnl1 bind in close proximity to the same 3' UTRs, and exert opposing effects on mRNA stability. a) A Venn diagram showing the expected and observed overlap between CELF1 and Mbnl1 3' UTR targets in myoblasts. The observed overlap is ∼3 times larger than expected (analysis controlled for gene expression). b) Expression change following CELF1 induction in muscle (7 d versus control) for transcripts grouped by number of Mbnl1 and CELF1 CLIP clusters in the 3' UTR. CLIP clusters for Mbnl1 were derived from myoblast, and for CELF1 from muscle. c) The probability density function (PDF) of the distribution of distances between CELF1 CLIP clusters in muscle and Mbnl1 CLIP clusters in myoblasts, in 3' UTRs with binding for both proteins. Distances for true binding sites are shown in blue, and for randomly placed binding sites in black. Statistical significance was assessed by modified KS test, and the distribution of distances in shuffled controls is shown in gray. d) Expression change following CELF1 induction in muscle (7 d versus control) for transcripts with exactly 1 Mbnl1 CLIP cluster and exactly 1 CELF1 CLIP cluster. Genes were grouped according to whether the distance between motifs is less than or greater than 50 base pairs. Also see Figure S5 and Table S5.

To assess possible functional connections between CELF and Mbnl in the cytoplasm, we assessed gene expression changes for sets of genes grouped by extent of CELF and Mbnl binding. Expression changes following *CELF1* induction exhibited a strong dependence on the ratio of CELF1 to Mbnl1 binding sites: greater CELF1 binding was associated with strong (up to ∼2-fold) down-regulation, messages with similar numbers of CELF1 and Mbnl1 clusters exhibited little change, and those with greater Mbnl1 binding were upregulated (Fig. 4b). These data suggest that Mbnl binding can protect a message from CELF-directed decay. The striking pattern in which each additional CELF binding site conferred reduced expression, while each additional Mbnl binding site conferred increased expression suggest that Mbnl1 and CELF1 compete for co-bound mRNAs to target them for different fates. Genes that were down-regulated upon CELF induction, are bound by both CELF and MBNL, and changed their localization upon MBNL depletion in the expected direction (away from membrane and towards the insoluble compartment) included *Cpe*, *Igfbp5*, *Kcnj2* and *Sobp* (see Discussion).

Among mRNAs not detectibly bound by CELF, the up-regulation of mRNAs bound more strongly by Mbnl relative to less-bound messages was unexpected. One possibility is that this signal derives from CELF1 CLIP false negatives (i.e. messages that are bound by CELF1 but which failed to be effectively detected by CLIP-seq). Another possibility is that CELF proteins may exert a general non-specific or low-specificity mRNA decay-promoting activity at high levels, and that Mbnl binding may protect against this effect.

To further explore the relationship between CELF1 and Mbnl1 proteins, we analyzed the relative locations of CELF1 and Mbnl1 binding sites in 3' UTRs. We found that sites bound by CELF1 and Mbnl1 tended to be closer together than expected, relative to randomly placed clusters (Fig. 4c; p < 1.8e-55 by modified Kolmogorov-Smirnov (KS) test). This proximity effect remained highly significant when performing additional analyses that preserved binding site locations but permuted protein identity, which controls for variations in accessibility along mRNAs (Fig. S5a).

To address whether functional antagonism between Mbnl1 and CELF1 might depend on binding site proximity, we examined the effects of CELF1 induction on messages with proximal (≤ 50 nt) or distal (> 50 nt) pairs of CELF1 and Mbnl1 binding sites. Notably, the presence of a distal Mbnl1 site was sufficient to abrogate the down-regulation normally associated with presence of a CELF1 site, but the presence of a proximal Mbnl1 site had no such protective effect (Fig. 4d). These effects extended to messages containing multiple CELF1 sites. Comparing sets of mRNAs with the same distributions of CELF1 site counts, and exactly one Mbnl1 site, those with a greater number of proximal CELF1-Mbnl1 pairs were more strongly downregulated following *CELF1* induction (Fig. S5b). These observations can be interpreted under a model in which CELF over-expression can displace proximal Mbnl1 (or Mbnl-associated protein complexes), but does not affect the ability of distally located Mbnl1 to antagonize CELF1-mediated mRNA decay.

mRNAs containing a proximal pair of Mbnl and CELF sites where protein displacement may occur are expected to be more responsive to developmental changes in CELF and Mbnl levels than messages containing distal pairs of sites. To explore this idea, we assessed developmental changes in expression of mRNAs grouped by binding site count and the presence of proximal or exclusively distal CELF/Mbnl site pairs (Fig. 4e). On average, messages were moderately de-repressed in proportion to the number of CELF sites present, and were repressed when Mbnl sites exceeded CELF sites, consistent with the established effect of CELF on mRNA decay and the protective effects of Mbnl binding observed above. However, messages with proximal CELF/Mbnl pairs were de-repressed more strongly than those containing exclusively distal pairs, supporting a model in which proximal pairs confer enhanced responsiveness to changes in the relative levels of these factors, e.g., as a result of direct physical displacement of Mbnl1 by CELF1 or direct interference with its activity.

### CELF1-mediated regulation of alternative 3' UTR isoforms in development

Many mammalian genes end with multiple polyadenylation sites (PAS), yielding “tandem UTR” isoforms, differing in the length of their 3' UTRs. We observed many instances of altered 3' UTR isoform abundance following CELF1 induction in heart and muscle. For example, the Cornichon Family AMPA Receptor Auxiliary Protein 4 (*Cnih4*) gene contains at least two “tandem” alternative PAS that are used in these tissues. The relative abundance of the longer isoform was ∼85% and ∼75% in adult heart and muscle, respectively, but decreased monotonically over the CELF1 induction time course to ∼30% and 40%, respectively (Fig. 5a). Numerous CELF1 CLIP clusters are located within the extension region of the 3' UTR (between the two PAS), suggesting that decreased abundance of the longer isoform results from CELF1-mediated destabilization of this isoform. Our genome-wide analysis revealed that 57 tandem UTR events contained at least 3 CELF1 clusters in the extension region and zero clusters within the core (shared) region; of these, ∼75% exhibited decreased abundance of the longer isoform relative to the shorter isoform following CELF1 induction. Across other combinations of binding sites, a general trend was observed with greater relative decline in distal isoform expression associated with greater relative abundance of CELF1 clusters in extension versus core regions (Fig. 5b).

In another class of genes, a combination of alternative splicing and PAS usage gives rise to alternative last exons (ALEs) (Fig. 5c). These distinct 3' UTRs provide natural “reporters” for the function of the regulatory elements they contain. For over 1000 genes, we observed a monotonic change in the relative expression of ALE isoforms following CELF1 induction in muscle. Within this set, the ALE isoform exhibiting greater CELF1 binding tended to be downregulated, and the magnitude of this bias was strongest for pairs with greatest differential binding (Fig. 5c). To avoid complications related to possible regulation of splicing by CELF1, UTR pairs lacking CELF1 binding within 500 nt downstream of each stop codon were analyzed. A similar bias was observed (Fig. S6), suggesting that changes in ALE abundance are primarily caused by differential regulation of message stability by CELF1. These results and those observed for tandem UTRs provide further evidence of widespread CELF1-mediated mRNA decay and identify many examples of isoform-specific regulation.

The dramatic changes in CELF1 levels that occur during heart development led us to ask whether these 3' UTR isoforms are often developmentally regulated. Comparing changes in 3' UTR isoform abundance during heart development to changes following CELF1 induction in heart, we observed a strong negative correlation (R_Sp_ = –0.52) (Fig. 5d). Therefore, as observed at the level of splicing, *CELF1* induction tends to reverse isoform changes that occur during normal heart development. This negative correlation held whether analyzing tandem UTRs or ALEs, and whether analyzing CELF1 induction in heart or skeletal muscle (Fig. 5e). The hundreds of 3' UTR isoforms involved in these patterns suggests that developmental reductions in CELF1 activity underlie a substantial fraction of 3' UTR isoform changes that occur during heart development.

**Figure 5.**
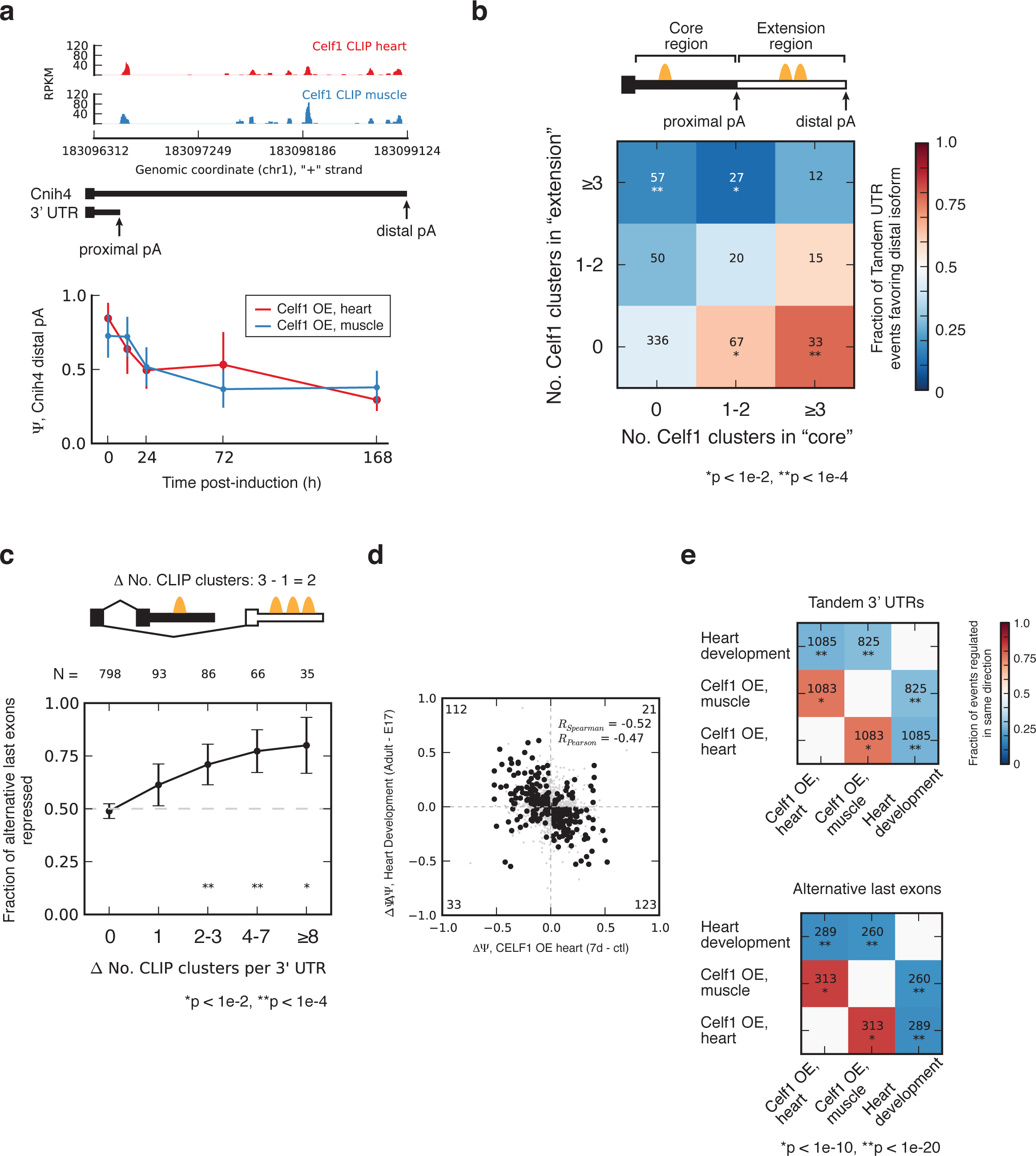
CELF1 regulates abundance of bound alternative 3' UTRs, reversing developmental changes. a) CELF1 CLIP-seq density in 3' UTR of Cnih4 gene, which has two alternative cleavage/polyadenylation sites (PAS) whose relative abundance changes following CELF1 over-expression and during heart development. b) CELF1 regulates tandem 3' UTR events in a manner that is dependent on the number of binding sites in the core and/or extension region of the 3' UTR. The heatmap shows the fraction of tandem UTR events biased towards usage of the proximal PAS following CELF1 over-expression, for events grouped by the number of CLIP clusters in the proximal or distal region of the 3' UTR. c) CELF1 regulates alternative last exon (ALE) expression in a manner that is dependent on the number of binding sites in the competing 3' UTRs. The fraction of ALEs that are significantly repressed following CELF1 over-expression, grouped by density in repressed 3' UTR minus density in enhanced 3' UTR. d) Changes in ALE usage during heart development inversely correlate with those that occur in response to CELF1 over-expression in heart. Correlation coefficients shown for events meeting a minimum Z-score threshold (heart development: 1.4, CELF1 OE: 1.8). As in Fig. 1c. e) Spearman correlation coefficients are displayed in heat map format for change in ALE usage (top) or change in tandem 3' UTR usage (bottom) for pairwise comparisons of isoform changes during heart development, CELF1 over-expression in heart, and CELF1 over-expression in muscle. Also see Figure S6.

## Discussion

The relationship between CELF and Mbnl proteins is important both in development and in the context of neuromuscular disease, particularly DM. Throughout mouse heart development, CELF protein levels decrease and Mbnl levels increase (Kalsotra et al. 2008); in DM, Mbnl proteins are sequestered by expanded CUG or CCUG repeats, and CELF proteins are stabilized via hyperphosphorylation by PKC and derepressed as a result of reduced miRNA expression (Timchenko et al., 2004; Kuyumcu-Martinez et al., 2007; Kalsotra et al., 2010). The nature of the relationship between these RNA binding proteins in normal physiology and development has been long-standing question in the field.

We observed a strong pattern of antagonistic regulation of mRNA levels by CELF and Mbnl associated with binding to 3' UTR regions. It is well established that CELF1 can recruit deadenylases to 3' UTRs and destabilize mRNAs (Vlasova et al., 2008), and recently we uncovered a global role for Mbnls in localizing mRNAs to membrane destinations for localized translation (Wang et al., 2012). These observations suggest a “Cytoplasmic Competition” (CC) model in which Mbnl and CELF proteins specify different cellular outcomes for mRNAs, and compete with one another to determine the fates and stabilities of specific mRNAs that contain binding motifs for both factors (Fig. 6a). Changes in localization often impact mRNA stability (Walters and Parker, 2014). Our data, e.g., the striking pattern shown in Figure 4b, can be most simply explained if messages tend to have a basal decay rate in the cytoplasm, which is generally accelerated by CELF1 induction and specifically enhanced by direct CELF1 binding, but have a lower decay rate when localized to membranes. Under this model, the localizations and half-lives of a over one thousand messages bound by both factors are determined by the outcome of a competition between the activities of Mbnl and CELF proteins.

**Figure 6.**
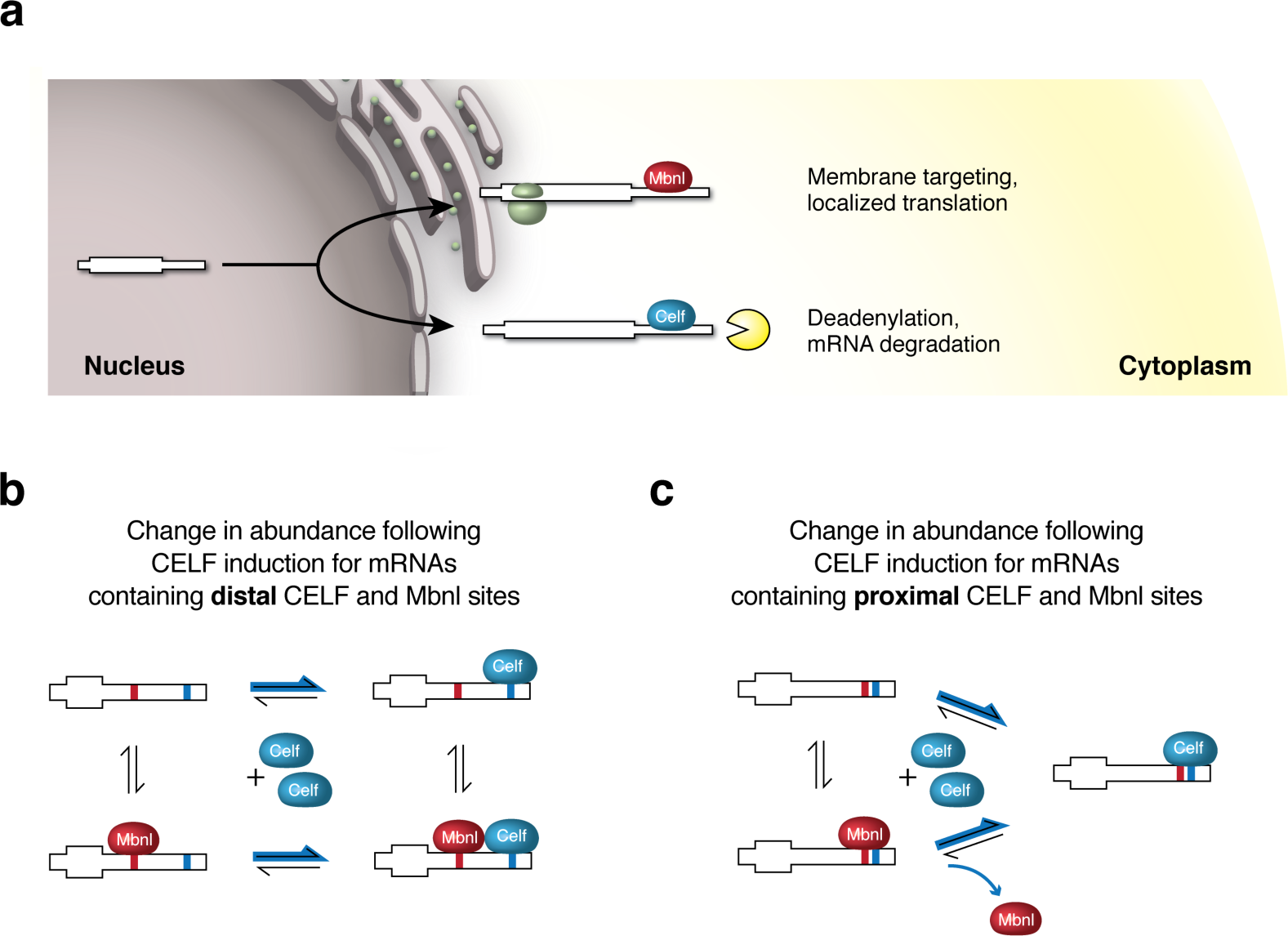
Cytoplasmic competition model of effects of CELF and Mbnl proteins on cytoplasmic mRNA fates. a) CELF1 binding to 3' UTRs promotes mRNA deadenylation and decay (Vlasova and Bohjanen, 2008), while Mbnl1 binding to 3' UTRs promotes localization to membrane compartments (Wang et al., 2012). In the cytoplasmic competition model, binding of both proteins to the same mRNA result in a tug-of-war, with Mbnl-directed localization promoting stabilization and CELF binding promoting decay. b) For mRNAs containing distal Mbnl and CELF binding sites, CELF1 over-expression shifts the equilibrium toward CELF-bound and doubly-bound mRNAs as indicated by the blue arrows. c) For mRNAs containing proximal Mbnl and CELF binding sites, CELF over-expression causes displacement of Mbnl (as illustrated), or of Mbnl-dependent complexes involved in mRNA localization/stabilization (not shown), effectively shifting the equilibrium toward mRNAs exclusively bound by, or controlled by, CELF (blue arrows).

Messages bound by both factors whose expression responded to CELF1 induction and whose localization changed in response to Mbnl depletion (Results) included *Cpe*, involved in insulin processing, and *Igfbp5*, which regulates insulin-responsive growth factor. Other messages with this pattern of binding, expression and localization change included *Kcnj2*, a cardiac inward rectifier potassium channel whose mutation is associated with a syndrome that involves cardiac arrhythmia (Andelfinger et al. 2002), and Sine Oculis binding protein homolog *Sobp*, mutations of which have been linked to intellectual disability (Birk et al., 2010). Thus, genes that represent competing cytoplasmic targets of CELF1 and Mbnl may contribute to symptoms like insulin resistance, cardiac arrhythmia, and mental retardation that are observed in DM.

Downregulation of the *Clcn1* gene, which encodes the major voltage-gated chloride channel that controls the membrane excitability of skeletal muscle, causes the myotonia that is characteristic of DM (Charlet et al., 2002; Mankodi et al., 2002). This downregulation is thought to result primarily from aberrant inclusion of exon 7a, due to loss of Mbnl activity, which produces a nonfunctional *Clcn1* mRNA (Wheeler et al., 2007). Here, we showed that *Clcn1* mRNA levels decreased monotonically following CELF1 over-expression in both muscle and heart, and that CELF1 binds to the *Clcn1* 3' UTR (Fig. 3c). These observations suggest that CELF1 may downregulate the stability of *Clcn1* mRNA, independent of splicing regulation. Therefore, the up-regulation of CELF activity that often accompanies Mbnl sequestration in DM skeletal muscle may destabilize the *Clcn1* mRNA and exacerbate myotonia symptoms.

The distance between CELF and Mbnl sites on a message appears to play an important role in the outcome of competition between these factors: close spacing of CELF1 sites near Mbnl sites (less than ∼50 nt apart) appears to override the protective effect normally conferred by Mbnl1 binding (Fig. 4d, 6b). This effect might result either from direct occlusion of Mbnl1 binding, or by antagonism of the binding or function of Mbnl1-dependent protein complexes. The functional antagonism between the cytoplasmic activities of Mbnl and CELF proteins uncovered here is likely to contribute to the robustness of developmental changes in mRNA stability and localization, particularly for mRNAs bound by both factors (Fig. 4e). These mRNAs may be particularly susceptible to mis-regulation in DM, perhaps contributing to pathogenesis.

We identified several dozen exons exhibiting antagonistic splicing regulation, many of which are developmentally regulated. These exons tend to preserve reading frame and to reside in genes affecting functions involved in development and cell differentiation (see Results). Human homologs of this set of antagonistically regulated exons may be mis-regulated in DM to a particularly strong extent, as increased levels of CELF proteins may exacerbate splicing changes that result from Mbnl sequestration. The hypothesis that CELF binding to exons tends to repress splicing, suggested by the genome-wide analyses, was confirmed in a splicing reporter system, establishing a rule to predict CELF-depending splicing regulation.

CELF1 function intersects with RNA processing in a different way in its impact on the stability of alternative mRNA isoforms that differ in their 3' UTRs. We observed a large set of genes with developmentally regulated alternative last exons and tandem UTR isoforms that differentially respond to CELF induction. Regulation of some of these isoforms by CELF1 may impact encoded protein functions by changing the abundance of protein isoforms differing in their C termini, while regulation of others may alter mRNA localization or translation or other UTR-dependent properties during development and other physiological situations in which CELF activity changes.

## Methods

### Heart Inducible CELF2 mouse

N-terminal Flag-tagged human CELF2 was expressed from the transgene previously used for CELF1 (Koshelev et al., 2010). TRECUGBP2 transgenic mice were generated by standard techniques and maintained on an FVB background. All reported TRECUGBP2/MHCrtTA bitransgenic mice were the F1 progeny from TRECUGBP2 x MHCrtTA (FVB/N-Tg(Myh6-rtTA)1Jam) matings. TRECUGBP2/MHCrtTA bitransgenic mice induced using doxycycline did not show evidence of ECG, echocardiography abnormalities or abnormal heart size or morphology after 8 weeks of induced expression of CELF2.

### RNA-seq Library Preparation and Sequencing

CELF1 and CELF2 were induced in heart and muscle by feeding mice 2 g/kg doxycycline for 12 hours, 24 hours, 72 hours, or 7 days, in the relevant transgenic animal. For CELF1 induction in heart, mice with myosin heavy chain promoter-driven reverse tetracycline transactivator (rtTA) and tet-inducible, N-terminal Flag-tagged human CELF1 LYLQ isoform were used. For CELF2 induction in heart, mice with the myosin heavy chain promoter-driven rtTA and tet-inducible human CELF2 were used. Hearts were harvested, atria removed, and ventricles frozen in liquid nitrogen. For CELF1 induction in muscle, mice with the rat myosin light chain 1/3 promoter/enhancer driving rtTA and tet-inducible, N-terminal Flag-tagged human CELF1 LYLQ isoform were used. The left and right gastrocnemius were isolated and frozen in liquid nitrogen. Control experiments for all time courses used mice with rtTA cassettes, but lacking the tet-inducible CELF cassettes; control mice were fed 2 g/kg doxycycline for 72 hours before tissue harvest. Total RNA was isolated in all cases using Trizol (Invitrogen), followed by RNeasy column with DNase treatment (Qiagen). Poly(A)+ RNA was prepared using Oligo dT Dynabeads (Invitrogen) and prepared for paired-end RNA sequencing (36 to 40 cycles by Illumina GA II).

### CLIP-seq library preparation and sequencing

CLIP was performed using 254 nm UV irradiation as previously described (Wang et al., 2009b), using heart tissue or muscle tissue of 16-week-old mice or cultured C2C12 mouse myoblasts.

Tissue was ground to a powder using liquid nitrogen-cooled mortar and pestle prior to UV irradiation. The dry powder was placed into a 10 cm^2^ tissue culture dish, sitting on ice, and crosslinked 3 x 400 mJ/cm^2^. In between each round of cross-linking, the dish was shaken from side to side, to redistribute the tissue powder and provide maximum opportunity for all tissue particles to be cross-linked. The tissue was then lysed in RIPA buffer (50 mM Tris-HCl pH 7.4, 150 mM NaCl, 0.1% sodium deoxycholate, 1% NP-40, 0.5% SDS). The lysate was treated with DNase and RNase I_f_ (NEB) for 10 minutes at 37 degrees, with dilutions of 1:10,000 and 1:50,000 providing optimal RNA fragment lengths for downstream purification. Immunoprecipitation was performed using the 3B1 antibody clone against CELF1 (Millipore) and protein A beads (Invitrogen). The beads were washed twice with RIPA and twice with RIPA containing 1M NaCl. The 3' adapter was pre-adenylated with ImpA (Hafner et al., 2008) and ligated while the RNA-protein complexes were on beads using T4 RNA ligase (Rnl2 truncation) in the absence of ATP (NEB). The complexes were run on SDS-PAGE gel, transferred to nitrocellulose, and isolated from membrane as previously described (Wang et al., 2009b). The protein was digested by proteinase K, and RNA was precipitated, and ligated to the 5' adapter using T4 RNA ligase (NEB). Two nucleotide-long barcode sequences were used at the 3' end of the 5' adapter, to allow for censoring of CLIP read PCR duplications. Ligated products were resolved by electrophoresis on TBE-urea polyacrylamide gels, isolated, and subjected to RT-PCR using Superscript III enzyme (Invitrogen) and Phusion polymerase (NEB). PCR primers contained indexes for sequence multiplexing, but for these experiments no samples were multiplexed. Adapter and primer sequences (5' to 3') were as follows:

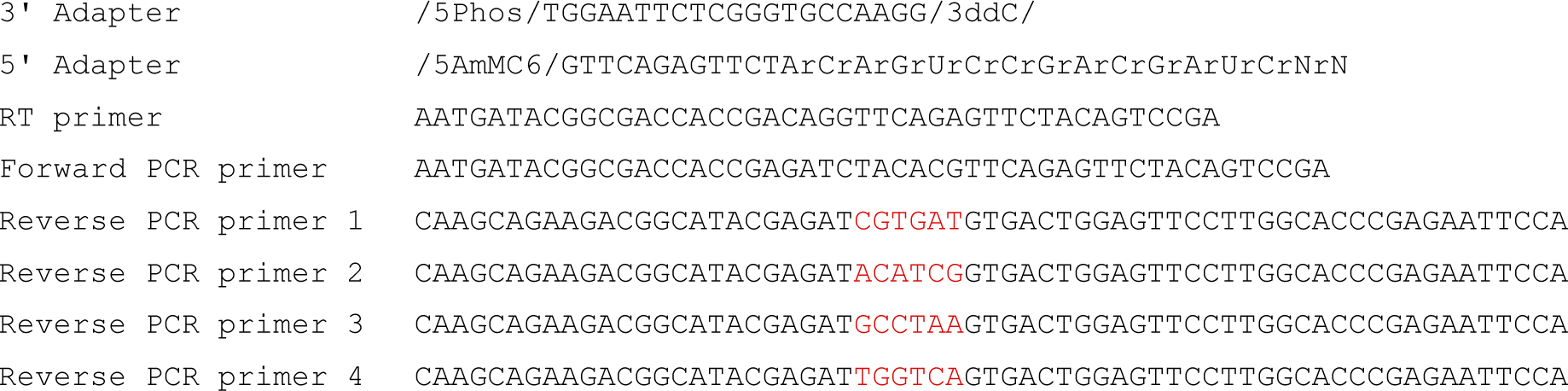

### RNA-seq and CLIP-seq read mapping, gene expression estimation

All read mapping was performed using Bowtie/Tophat (Langmead et al., 2009; Trapnell et al., 2009), mapping to mm9. Only uniquely mapping reads were used. CLIP reads were collapsed to remove identical sequences, adapter sequences were removed, and processed reads were then mapped separately for each CLIP read length. To estimate gene expression levels, the number of reads mapping to each kilobase of constitutive coding sequence of Refseq/Locuslink genes was counted, and divided by the number of reads (in millions) mapping uniquely to non-ribosomal and non-mitochondrial sequence, to obtain RPKM values. For purposes of the analyses performed in Fig. 4e, gene expression values in the heart development time course were normalized as described (Robinson and Oshlack, 2010), using parameters of M=0.3 and A=0.2.

### Estimation of isoform frequencies, calculation of MZ score

MISO (version 0.4.8) was used to estimate isoform frequencies for splicing events and alternative 3' UTR events, using a minimum of 20 reads per event and parameters of burn_in=500, lag=10, num_iters=5000, and num_chains=6. To identify splicing events that change monotonically over time, we ordered samples chronologically, and for each event compared all pairs of samples from different time points, tallying the number of comparisons representing significant increase or significant decrease in Y (at BF > 5). We calculated a quantity called d, the number of significant positive DY values (increases over time) minus the number of significant negative DY values. To assess statistical significance, we recalculated d after randomly permuting the sample labels. Repeating this process 100 times, we generated a null distribution, and derived the “monotonicity Z-score” (MZ = (d-m) / s), where m and s are the mean and standard deviation of the null distribution, respectively.

### Analysis of antagonistically regulated splicing events (Figs. 1g, 1h)

Splicing events regulated in response to each perturbation (CELF over-expression, Mbnl1 KO, or heart development) were enumerated, and the number of events regulated among each pair of perturbations was counted (this was the “observed” overlap). To compute the “expected” overlap, we assumed independence, e.g. the fraction of events regulated in both perturbations equals the fraction of events regulated in the first perturbation multiplied by the fraction of events regulated in the second perturbation. Significance of the bias in direction of regulation (Figure 1h) was assessed by binomial test, assuming a null hypothesis frequency of 0.5.

### Correlation of CLIP binding density across the transcriptome (Fig. 2b)

CLIP read density was computed in 5 nucleotide-long windows across 3' UTRs in the transcriptome for genes highly expressed in whole brain, heart, muscle, and C2C12 mouse myoblasts (>100 RPKM) exhibiting high CLIP coverage (>100 tags per UTR). Pearson correlation coefficients were computed for these densities between CLIP libraries performed in different samples and for different proteins.

### Identification of CLIP clusters

CLIP tags were first collapsed to remove redundant sequences, and trimmed of adapters. These sequences were mapped to genome and a database of splice junctions using Bowtie. To identify CLIP clusters lying within genic regions, gene boundaries were first defined using Refseq, Ensembl, and UCSC tables. For each window of 30 nucleotides covered by at least one CLIP tag, a test was performed to assess CLIP density in the window exceeded that which is predicted by a simple Poisson model which accounts for gene expression and pre-mRNA length (Yeo et al., 2009).

### CLIP cluster motif analysis (Fig. 2c, Fig. S3b)

Pentamers occurring in CELF1 CLIP clusters from heart were counted and compared to those found in randomly selected clusters within the same 3' UTRs (Fig. 2c) or whole genes (Fig. S3b). This procedure was repeated 100 times to derive a Z-score for each motif, where the Z-score was defined as the number of standard deviations away from the mean. The 5 most highly enriched pentamers are highlighted.

### Conservation analysis for cassette exons and 3’ UTRs (Figs. 2d, 3b)

Conservation (30-way phastCons, UCSC genome browser) of cassette exons with CLIP clusters within 1 kilobase of each splice site or within the exon itself was compared to a control set of cassette exons found in similarly expressed genes. Conservation (30-way phastCons) of 3' UTRs with CLIP clusters less than 1 kilobase upstream and less than 500 bases downstream of constitutive transcript ends was compared to a control set of 3' UTRs with similar length, in similarly expressed genes. For these analyses, CLIP clusters and gene expression values from heart were used.

### Crosslink-induced substitution analysis (Figs. 2e, S3c)

As in (Wang et al., 2012), p(CIS) was defined as the fraction of reads coverage a base supporting a base substitution differing from reference. Only positions with minimum read coverage above 10 were considered in this analysis. Information content of each base flanking frequently substituted cytosines was computed as 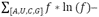 where *f* is the frequency of each base.

### Analysis of gene expression changes as a function of CELF CLIP clusters and/or microRNA seed matches (Fig. 3e-g)

Genes were grouped by the number of CELF CLIP clusters found in 3' UTRs (Fig. 3e) or microRNA seed matches (Fig. 3f), and the mean log2(fold-change) in expression level in each group was computed, relative to genes with no CLIP clusters or microRNA seeds, respectively. Significance was assessed by KS test. The abundance of microRNAs in heart (Fig. 3g) was derived from (Li et al., 2013).

### Analysis of binding targets shared by CELF1 and Mbnl1 (Fig. 4a)

Genes were binned by expression level in mouse myoblast (Wang et al., 2012), and within each bin, the number of genes with 3' UTRs containing CELF1 CLIP clusters only, Mbnl1 CLIP clusters only, or clusters for both proteins was counted (“observed” number). The “expected” number was computed by assuming independence. The ∼3-fold observed/expected value was obtained by taking the mean observed/expected value across all gene expression bins. In Figure 4a, CLIP data was derived for both CELF1 and Mbnl1 from myoblasts.

### Analysis of CELF1 and Mbnl1 binding locations within 3' UTRs (Figs. 4c-e, S5a-b)

To precisely assess the distance between CELF1 and Mbnl1 binding locations within 3' UTRs, we searched for known CELF1 and Mbnl1 binding motifs within each CLIP cluster for each protein. The motifs were derived from Bind-n-Seq data (Lambert et al., 2014). For Mbnl1, the motifs were: GCTT, CGCT, TGCT, GCGC, CCGC, CTGC, GCTA, ACGC, CGCA, AGCT, TTGC, and CAGC. For CELF1, the motifs were: TGTT, ATGT, TTGT, TGTC, GTGT, TGTA, GTTT, TGTG, GTCT, and TTTT. If none of these motifs was found within the cluster, the binding location was estimated to be the center of the CLIP cluster. The closest distance between each CELF1 and Mbnl1 motif derived from its respective cluster was recorded, and compared to randomly assigned clusters (Fig. 4c) or clusters whose identity was shuffled (Fig. S5a). Tests for significant differences in Figs. 4e-f and Fig. S5b were performed by modified KS test, where cumulative distributions were visually inspected to confirm a true shift in median values.

### Analysis of alternative 3' UTR isoforms (Figs. 5b-c, S6)

Abundance of alternative 3' UTR isoforms by MISO was performed in “multi-isoform” mode. However, analysis was restricted to those events with exactly 2 isoforms whose regulation is significant and monotonic (BF > 5, MZ MZ > 1.6 for Tandem UTRs, MZ > 1.5 for alternative last exons).

### Data Access

The RNA-seq and CLIP-seq data presented here have been submitted to GEO (accession number pending).

## Acknowledgements

We thank M. Swanson and members of the Burge lab for helpful comments.

## Supplementary Figure Legends

**Figure S1.** Gene expression levels as estimated by RNA-seq, for genes induced during CELF1 or CELF2 time courses. a) CELF1 mRNA expression in heart, at various time points following CELF1 induction in heart. b) CELF1 mRNA expression in muscle, at various time points following CELF1 induction in muscle. c) CELF2 mRNA expression in heart, at various time points following CELF2 induction in heart. d) CELF2-Flag fusion protein expression by Western (HRP-conjugated anti-Flag antibody, Sigma A8592), and total protein stain by Ponceau S for each heart sample. 90 ug of total protein from the apex of each heart was loaded.

**Figure S2.** Method for determining monotonicity of splicing regulation. a) Method for determining monotonicity score, MZ, of splicing changes. b) The number of splicing events regulated during heart development that are also responsive to CELF1 induction were counted, at various monotonicity Z-score (MZ) thresholds. The fraction of events at each threshold is also listed and represented by color in the heatmap. The combination of MZ thresholds used to create the scatter plot shown in Figure 1e is boxed in red.

**Figure S3.** CELF1 CLIP-seq in heart, muscle, and myoblasts. a) The proportion of reads mapping to 5’ UTRs, coding sequence, introns, and 3’ UTRs is shown as pie charts for CLIP-seq and RNA-seq from heart, muscle, and myoblasts. The ratio between these values is shown as bar plots, illustrating enrichment for introns and 3’ UTRs in CLIP reads relative to RNA-seq reads. b) Motif enrichment Z-scores for all pentamers occurring in CELF1 CLIP-seq clusters, relative to randomly selected genic regions, are shown for CELF1 CLIP clusters identified in heart, muscle, and myoblasts.

**Figure S4.** CELF1 binding is moderately associated with splicing regulation. Splicing events were binned by gene expression level (RPKM), and the observed fraction of splicing events with both binding and splicing regulation was plotted in black dots. Splicing regulation was defined using a minimum monotonicity score of 2, and binding between 1 kb upstream of the 3’ splice site and 1 kb downstream of the 5‘ splice site was counted. The expected fraction of events with both binding and regulation was estimated by assuming independence, and plotted in gray dots. A Fisher's Exact test was performed to assess whether the observed fractions were different than expected, at each gene expression level bin. The analysis was performed separately in heart and muscle, for splicing events with at least 1, 2, or 3 CELF1 CLIP clusters. There is a modest (∼10-40%) enrichment for splicing regulation of bound splicing events.

**Figure S5.** CELF1 and Mbnl1 binding sites are closer than expected. a) The distribution of observed and expected distances between the centers of CELF1 and Mbnl1 CLIP clusters. Plots on the left were generated using CELF1 CLIP clusters from heart, and on the right, CELF1 CLIP clusters from muscle. All Mbnl1 CLIP clusters were derived from myoblasts. Expected distances for plots in the top row were computed by reassigning the clusters to random locations within 3’ UTRs. For plots on the bottom, expected distances were computed by keeping the clusters in the same location, but randomly reassigning clusters as being bound by CELF1 or Mbnl1. b) Expression change following CELF1 induction in muscle (7 d versus control) for genes with 1 Mbnl1 CLIP cluster and more than 1 CELF1 CLIP cluster, grouped by the number of CELF1 sites < 50 nt away from the Mbnl1 site and normalized to genes with no CELF1 sites < 50 nt away from the Mbnl1 site.

**Figure S6.** Alternative last exon abundance as a function of difference in number of CELF1 CLIP clusters between each 3’ UTR. The fraction of ALEs that are significantly repressed following CELF1 over-expression, grouped by density in repressed 3' UTR minus density in enhanced 3' UTR. Analyses were performed using UTR pairs in which CELF1 binding is at least 100 nt (a), 300 nt (b), or 500 nt (c) away from each stop codon.

## Supplementary Table and Figure Summary

Table S1. Mice used for RNA-seq. Related to Figure 1. (Excel file)

Table S2. All splicing events detected in CELF1 and CELF2 OE time courses. Related to Figure 1. (Excel file)

Table S3. Splicing events regulated by both CELF and Mbnl, along with information about regulation during heart development. Related to Figure 1. (Excel file)

Table S4. Gene expression changes responsive to CELF induction. Related to Figure 3. (Excel file)

Table S5. CELF1 and Mbnl1 CLIP cluster locations within 3' UTRs. Related to Figure 4. (Excel file)

Figure S1. Gene expression levels as estimated by RNA-seq, for genes induced during CELF1 or CELF2 time courses. Related to Figure 1.

Figure S2. Method for determining monotonicity of splicing regulation. Related to Experimental Procedures and Figure 1.

Figure S3. CELF1 CLIP-seq in heart, muscle, and myoblasts. Related to Figure 2.

Figure S4. CELF1 binding is moderately associated with splicing regulation. Related to Figure 2. Figure S5. CELF1 and Mbnl1 binding sites are closer than expected. Related to Figure 4.

Figure S6. Alternative last exon abundance as a function of the difference in number of CELF1 CLIP clusters between each 3' UTR. Related to Figure 5.

